## Supplemental Figures with captions for "Seasonal Enhancement of the Viral Shunt Catalyzes a Subsurface Oxygen Maximum in the Sargasso Sea"

**Supplementary Figures**

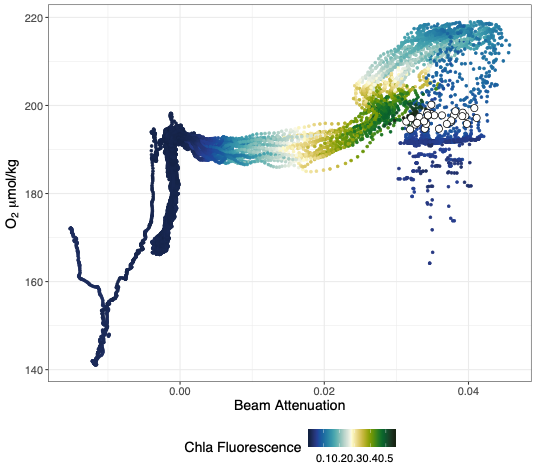
**Supplementary Figure 1. Correlation between oxygen level and beam attenuation, and chlorophyll *a* florescence.** Beam attenuation is a proxy for particle content. Each dot represents a CTD measurement from the 5-500 m depth profile during the October 2019 time series at BATS.

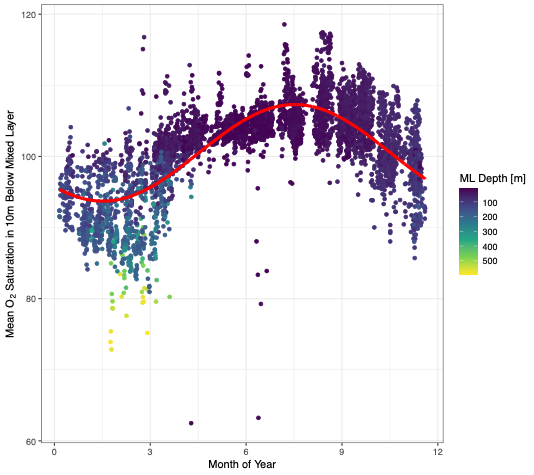

**Supplementary Figure 2. Seasonal cycle in sub-mixed layer oxygen dynamics at BATS.** Each point represents the mean oxygen saturation percentage for one of 5,054 casts from BATS cruises where optode data matched Winkler method bottle calibrations (see Methods). The x-axis is the month of the year in which the BATS cruise took place, the y-axis is the mean oxygen saturation between the mixed layer depth and 10 m below the mixed layer depth, with mixed layer depth determined by a change in potential density of 0.125 kg/m^3 from a reference pressure of 10 db. Point color indicates the calculated mixed layer depth of that cast. A nonlinear least squared fit of a sinusoidal regression was calculated and is shown as the red line. The model specification is: mean O_2_ saturation = a+b*sin(2pi*(c+decimal year)). Decimal year goes from 0 (Jan 1) to 1 (Dec 31). Estimated parameters with standard errors and p-values are: a=100.502, SE=0.0584, p<1e-10, b=-6.792, SE=0.0855, p<1e-10, c=0.123, SE=0.00186, p<1e-10.

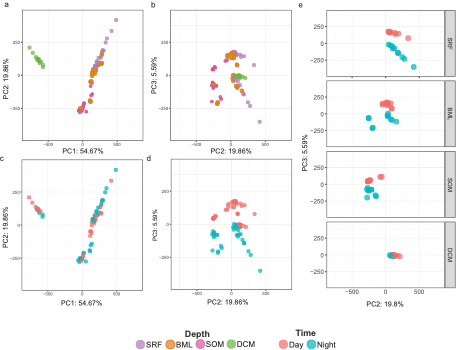

**Supplementary Figure 3.** **Principal components (PC) analysis of community metatranscriptomes show shifts in expression profiles with depth and time.** PCs were generated using normalized transcripts of all assembled genes across the co-assembly. a) PC1 against PC2 color coded by depth. b) PC2 against PC3 color coded by depth. c) PC1 against PC2 color coded by day (8:00, GMT-3) versus night (20:00, GMT-3). d) PC2 against PC3 color coded by day versus night. e) PC2 against PC3 color coded by time, plotting each depth separated into panels.

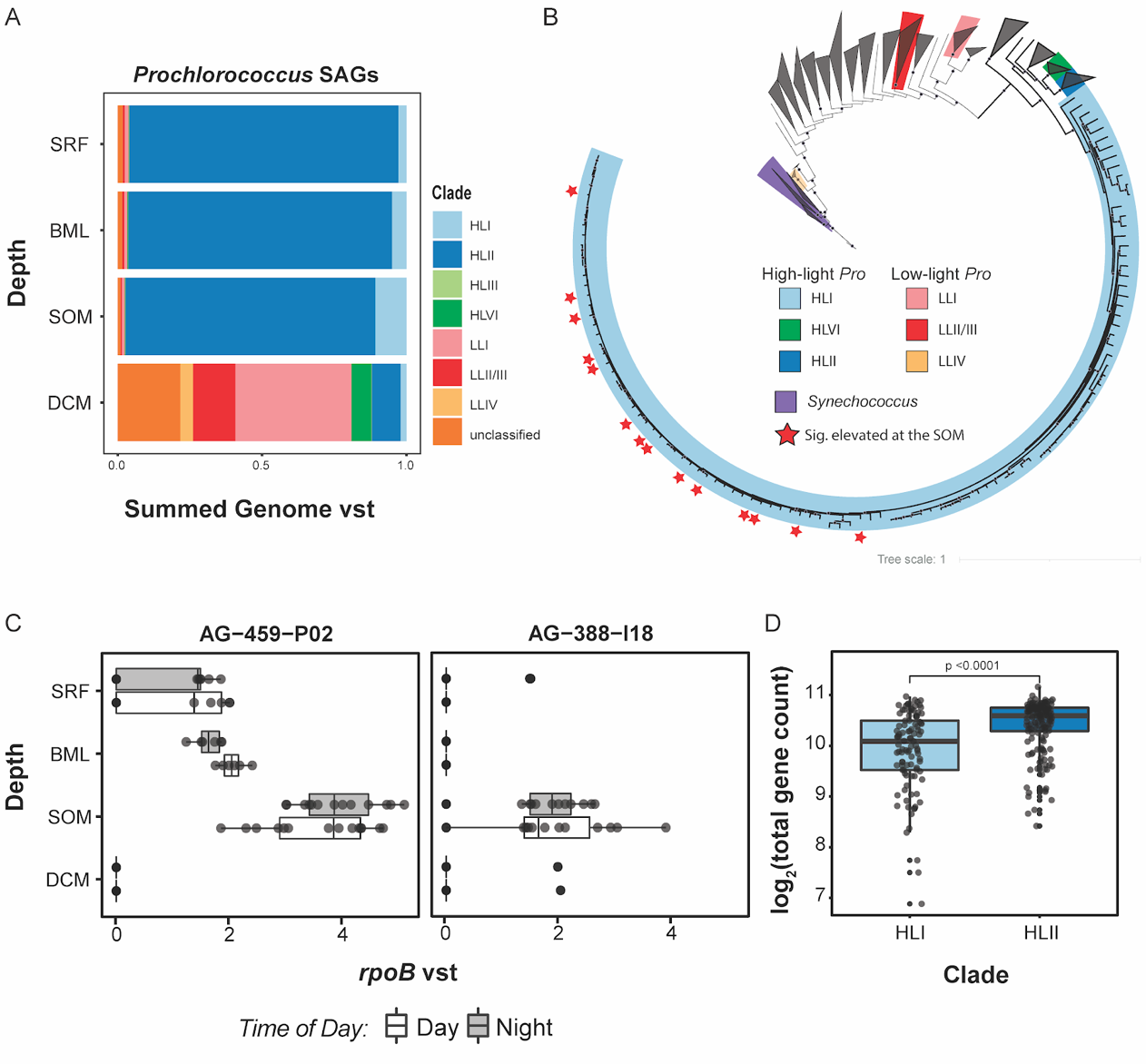

**Supplementary Figure 4. *Prochlorococcus* ecotype transcript representation across depth.** a) Stacked bar plot showing the distribution of transcripts assigned to single-celled amplified genomes (SAGS) from different *Prochlorococcus* ecotypes across depth. b) Maximum Likelihood phylogeny of cyanobacterial genomes. The branches are color coded by respective taxonomy. Stars indicate SAGs that had significantly elevated *rpoB* transcript abundances unique to the SOM. Because these all fell into the HLI clade, the other clades are collapsed for simplicity. SAGs and cyanobacterial genome were taken from Berube et al.^77^

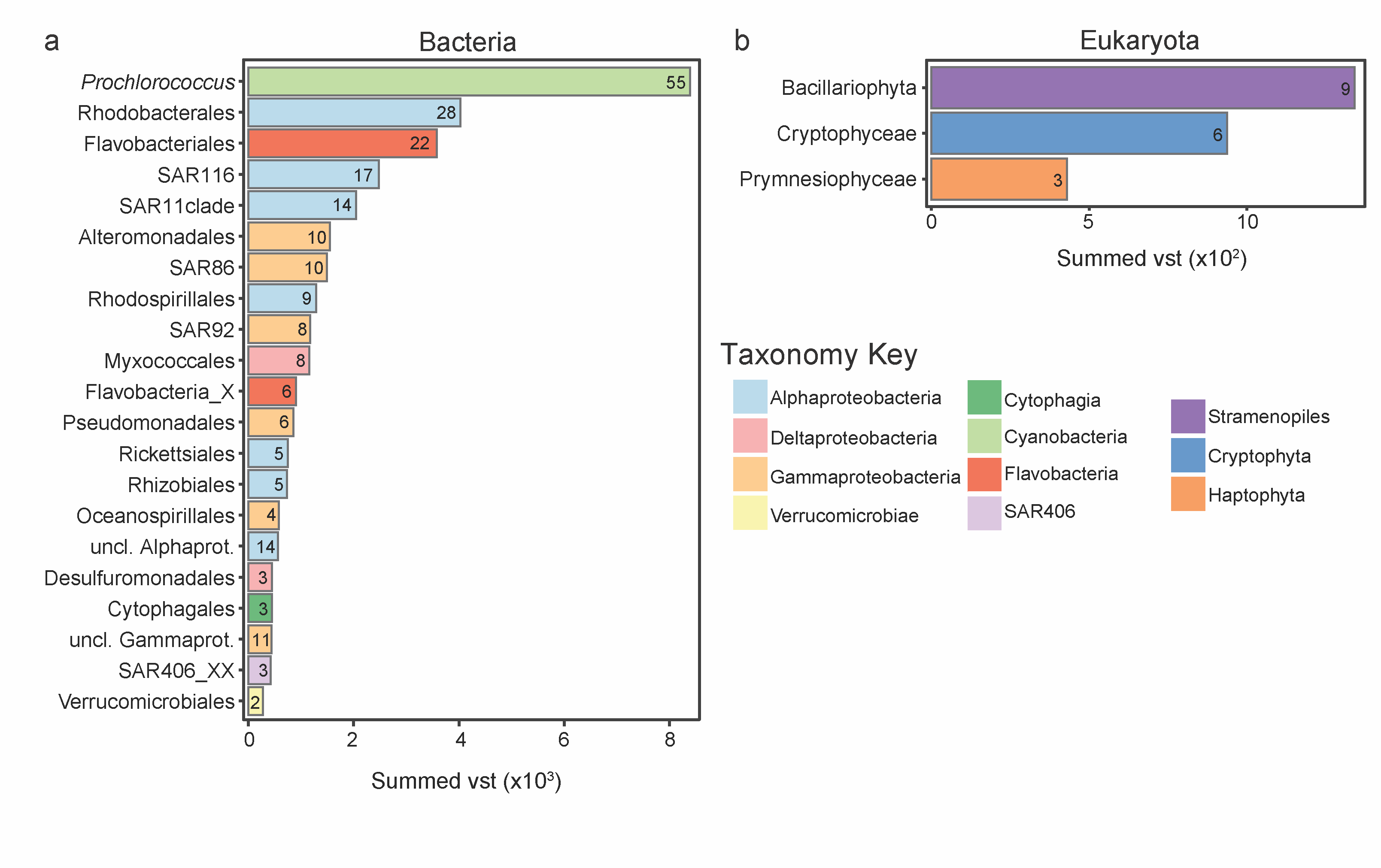

**Supplementary Figure 5. Taxonomic annotations of *rpoB/RPB1* harbored on individual contigs (*rpoB/RPB1* hit) with significantly increased transcripts at the SOM.** a) Summarized transcript abundances of top bacterial *rpoB/RPB1* hits with significantly elevated transcript abundances (BH-adjusted p≤0.1) specific to the SOM using order-level taxonomic annotation. *Prochlorococcus*-like *rpoB*s are annotated to the genus level. b) Summarized transcript abundances of top eukaryotic *RPB1* hits with significantly elevated transcript abundances (BH-adjusted p≤0.1) specific to the SOM using class-level taxonomic annotation. The number of individual *rpoB/RPB1* hits harbored on different contigs detected as significantly elevated at the SOM is shown within each bar, color coded by class-level information, for both panel a and b.

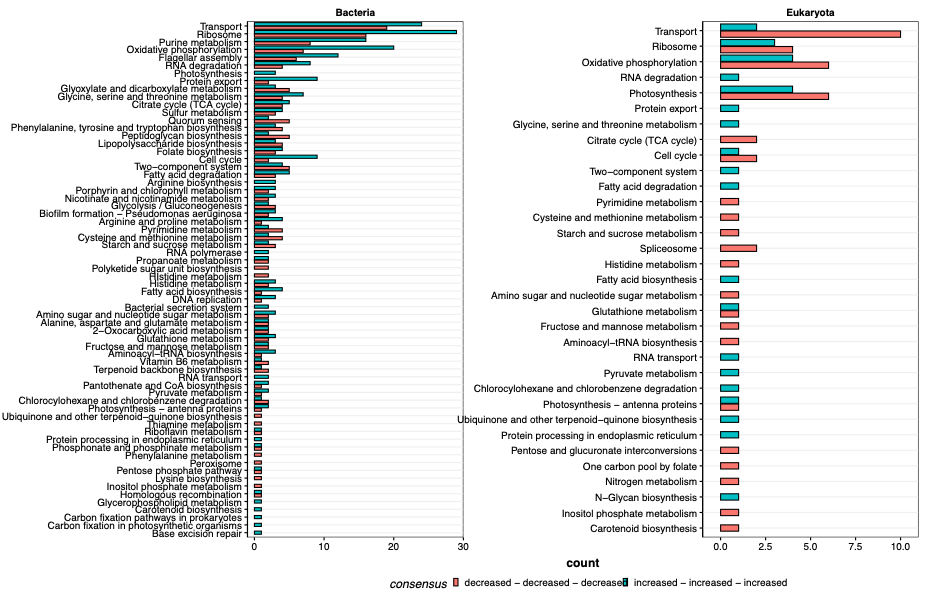

**Supplementary Figure 6. Expressed KEGG pathways either uniquely heightened or lowered at the SOM relative to the SRF, BML and DCM.** Top KEGG Orthology (KO) pathways assigned to KOs detected as significantly (BH-adjusted p≤0.1) heightened or lowered at the SOM for a) Bacteria and b) Eukaryotes. Red = heightened at the SOM, Blue = lowered at the SOM.

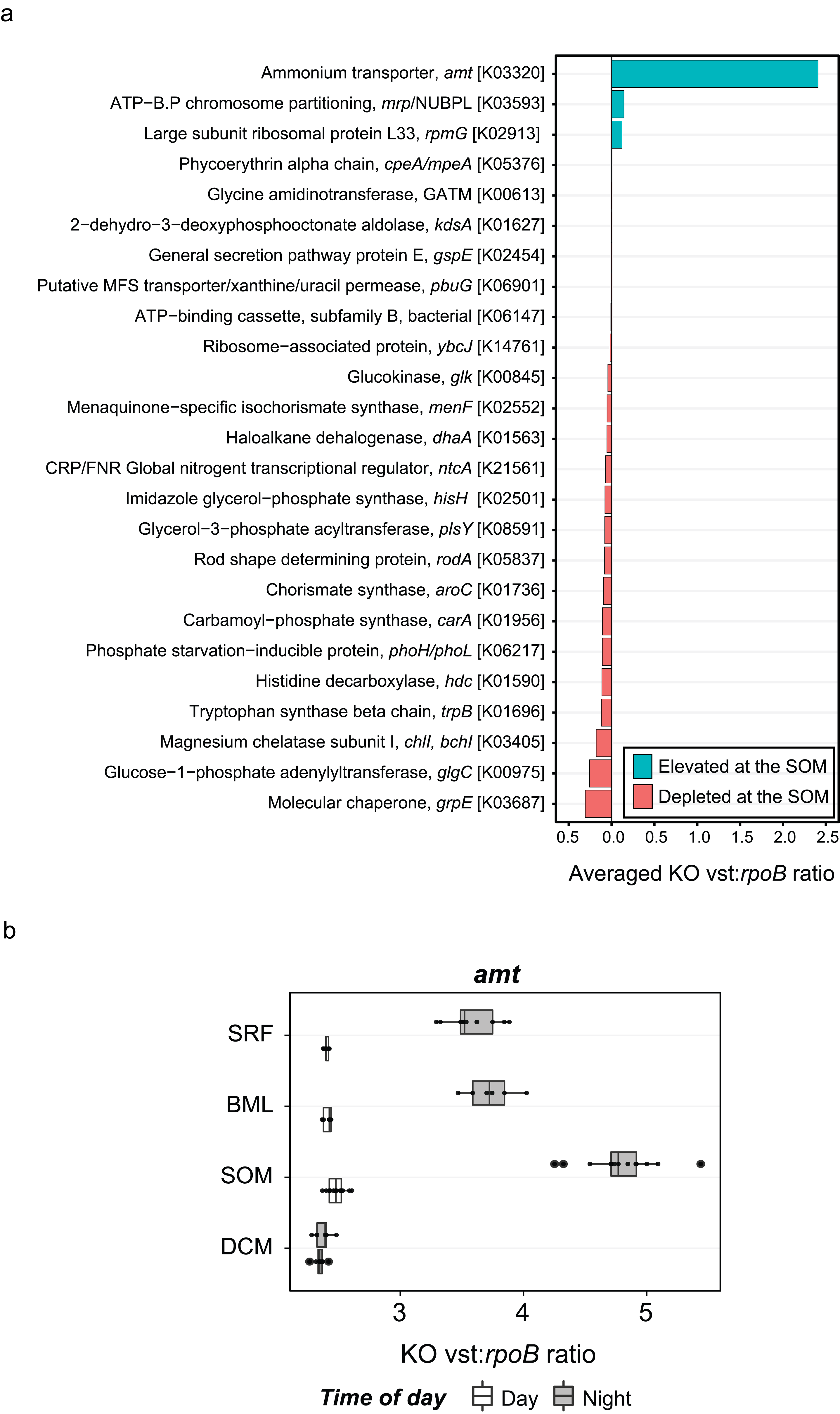

**Supplementary Figure. 7.** *Prochlorococcus-*specific genes with either uniquely heightened or lowered transcript values at the SOM. a) KOs assigned to *Prochlorococcus* with significantly (BH-adjusted p≤0.1) elevated or depleted transcript values at the SOM relative to the mixed layer and DCM, displayed as the ratio of the KO VST to bulk *Prochlorococcus rpoB* VST (VST ratio) within each sample, averaged across the entire dataset. b) Depth and time-related trends of the VST ratio of the *amt* gene (ammonium transporter gene, K03320).

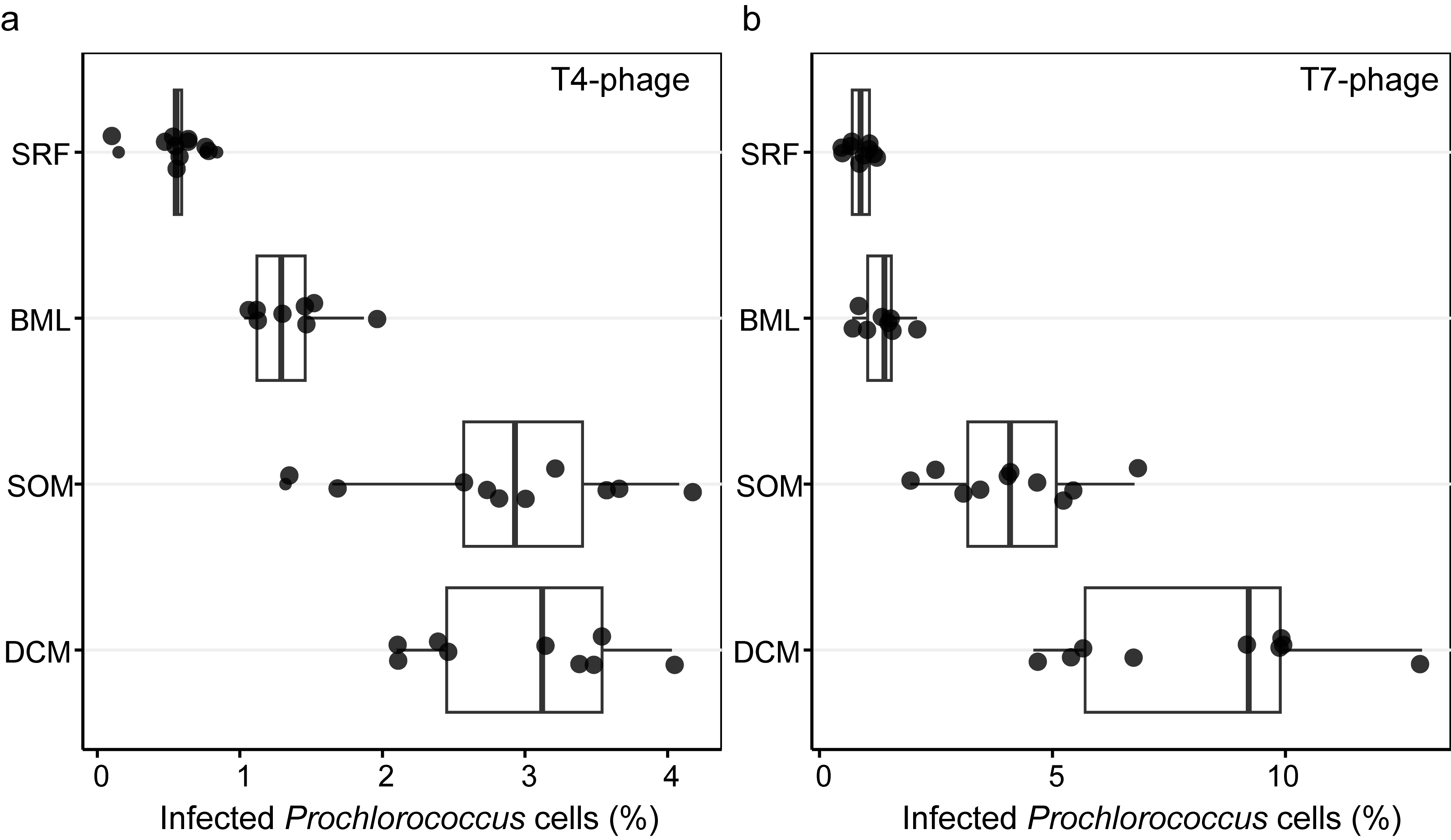

**Supplementary Figure 8**. **Extent of infection of *Prochlorococcus* cells by T4-like and T7-like cyanophages.** Percent infected *Prochlorococcus* cells by a) T4-like cyanophages and b) T7-like cyanophages. Infection was determined by the iPolony method quantifying the number of *Prochlorococcus* cells that have detected phage DNA, then calculating the percent of cells with phage DNA within the total number of cells interrogated.

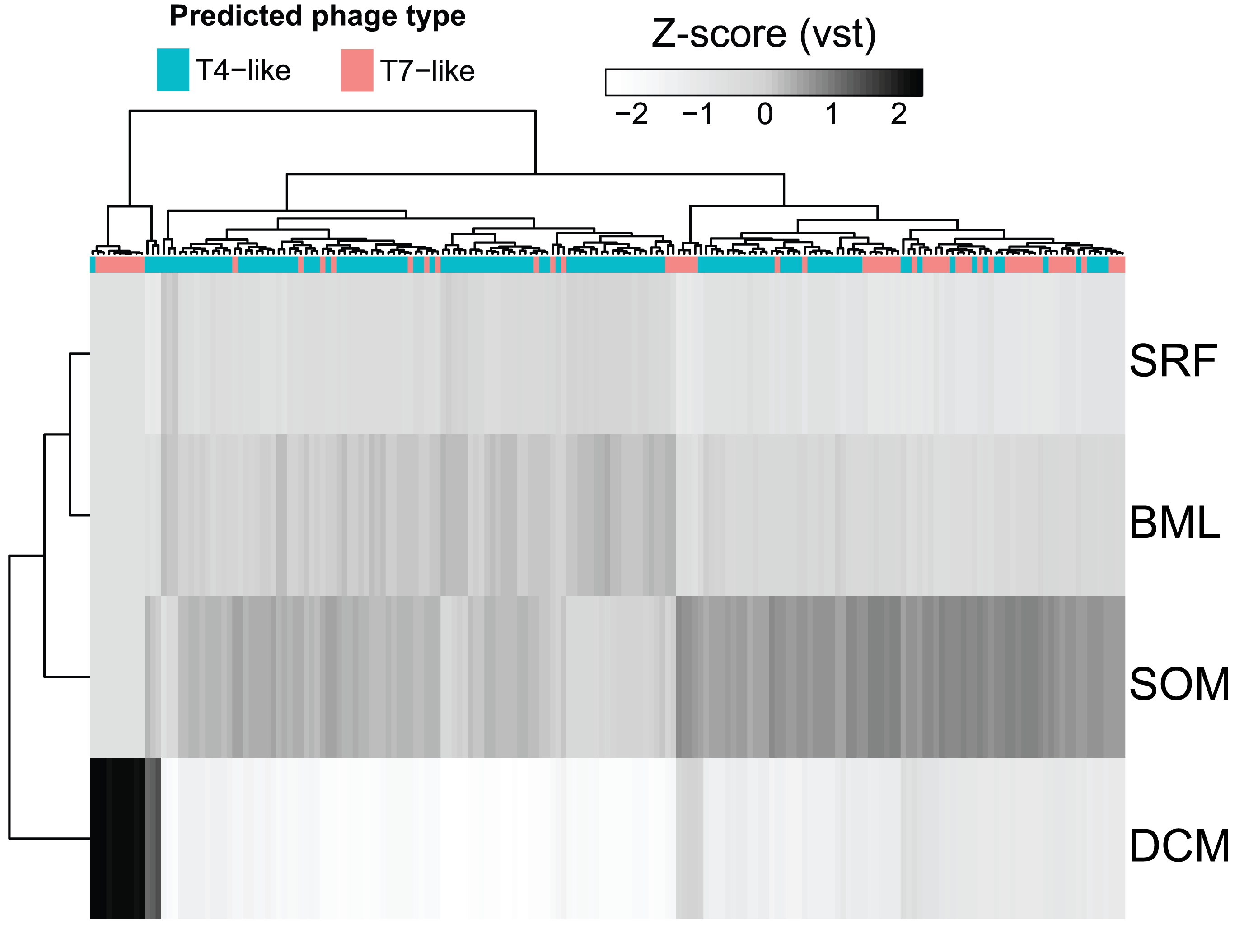

**Supplementary Figure 9. Expression patterns of all potential T7-like and T4-like phages broadly infecting prokaryotes detected in the viromics dataset.** Hierarchical clustering of normalized transcript abundance (VST) across entire vOTU scaffolds with significantly different transcript abundances as a function of depth. Values are averaged across depth for each vOTU. Putative taxonomy was determined by screening for T7-like and T4-like phages marker proteins on each scaffold.

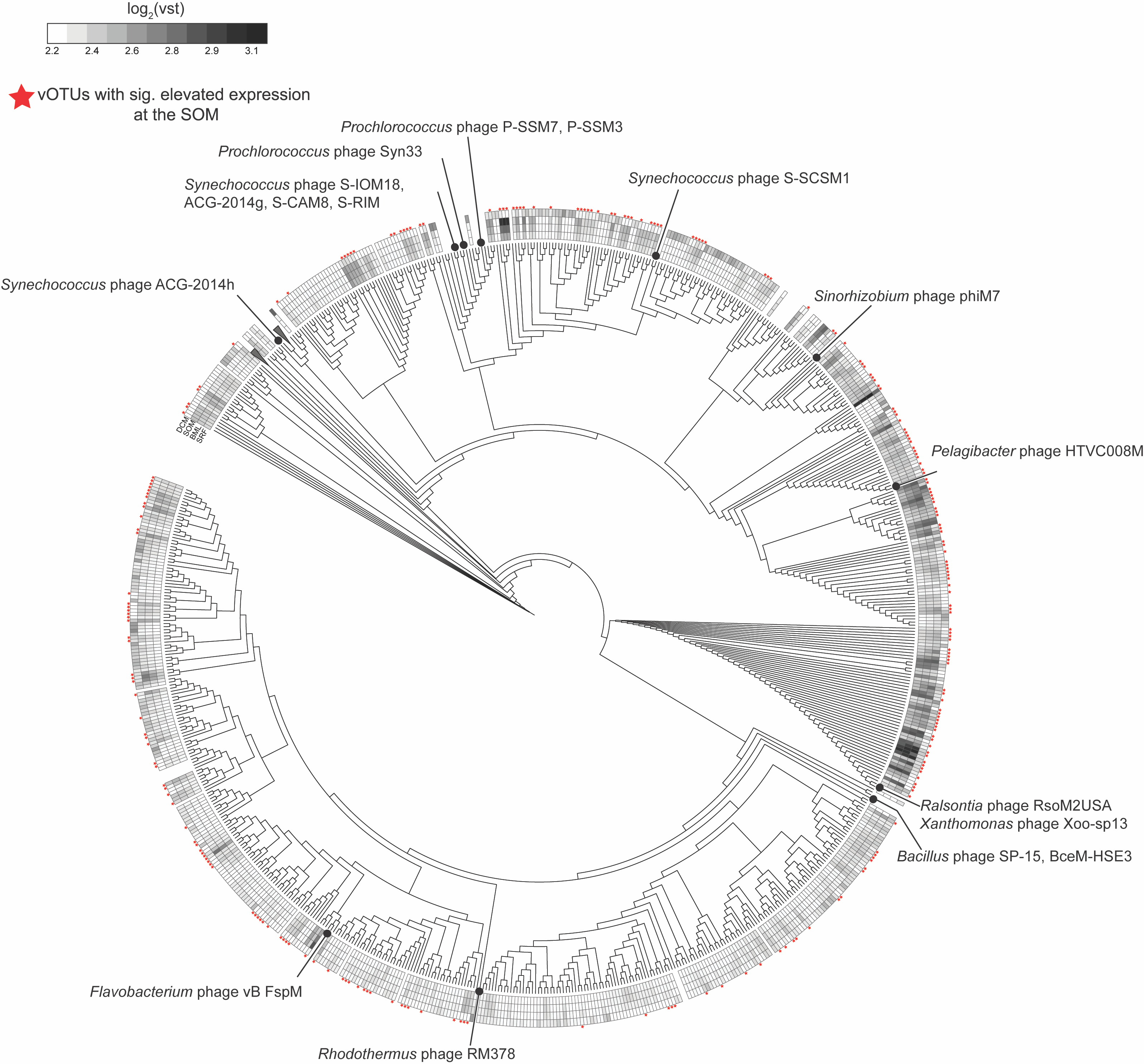

**Supplementary Figure 10. Expression of T4-like vOTUs from the viromes broadly infecting prokaryotes**. Phylogeny of T4-phage major capsid proteins (gp23) detected on vOTU scaffolds. Branches are color coded by their putative family-level assignment. The heatmap surrounding the tree shows depth-integrated log_2_(VST) values, from the SRF (inner ring) to the DCM (outer ring). vOTU gp23 transcripts with significantly elevated values at the SOM (p_adJ_ < 0.1) are indicated with a red star. References with assigned hosts used in the base tree are shown in their locations along the branches.

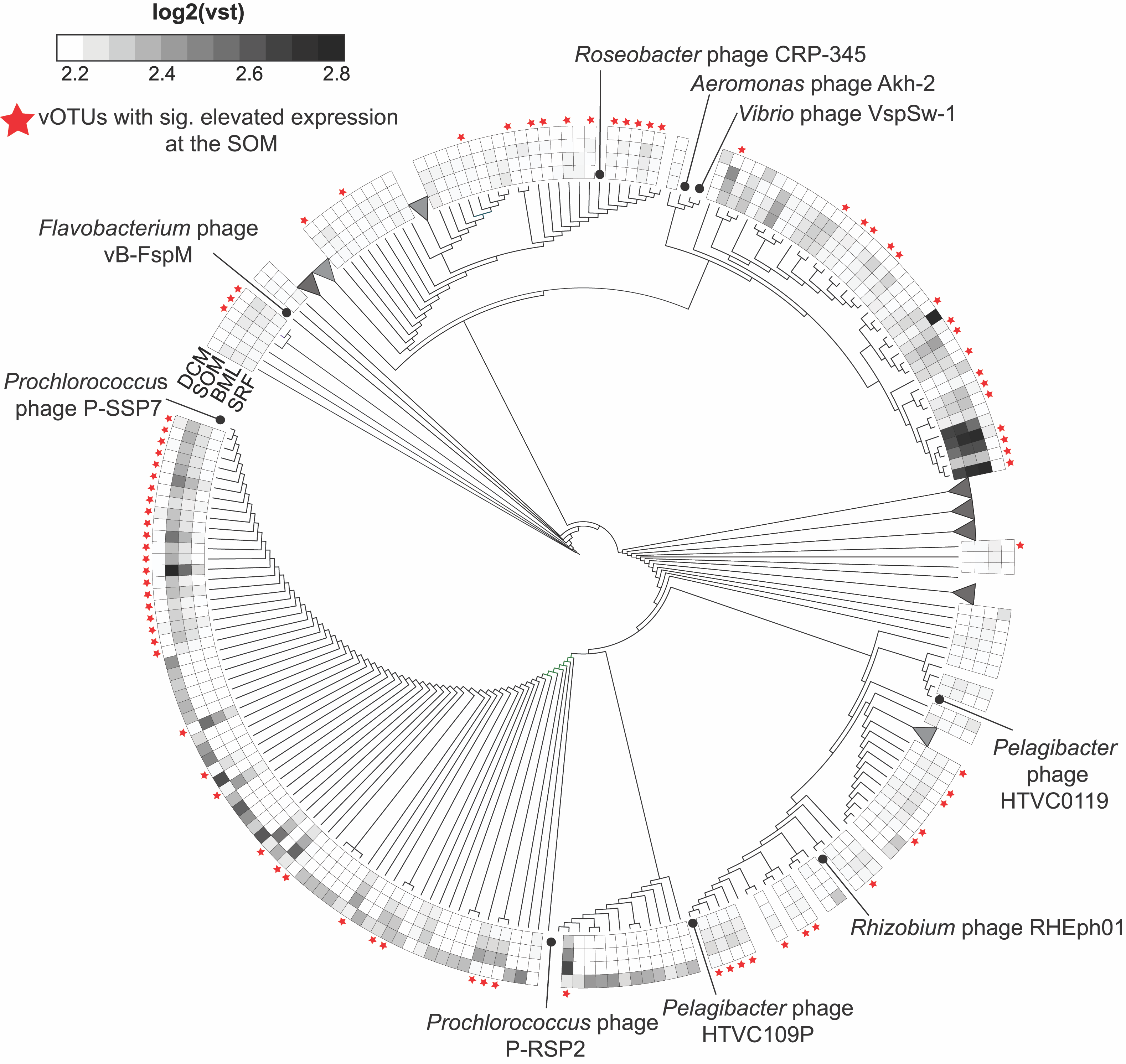

**Supplementary Figure 11**. Expression of T7-like vOTUs from the viromes broadly infecting prokaryotes. Phylogeny of DNA polymerase alpha-subunit (DNA Pol A) hallmark proteins detected on vOTU scaffolds. Branches are color coded by their putative family-level assignment. The heatmap surrounding the tree shows depth-integrated log_2_(VST) values, from the SRF (inner ring) to the DCM (outer ring). vOTU DNA Pol A transcripts with significantly (p_adJ_ < 0.1) elevated values at the SOM are indicated with a red star. References used in the base tree are shown in their locations along the branches.

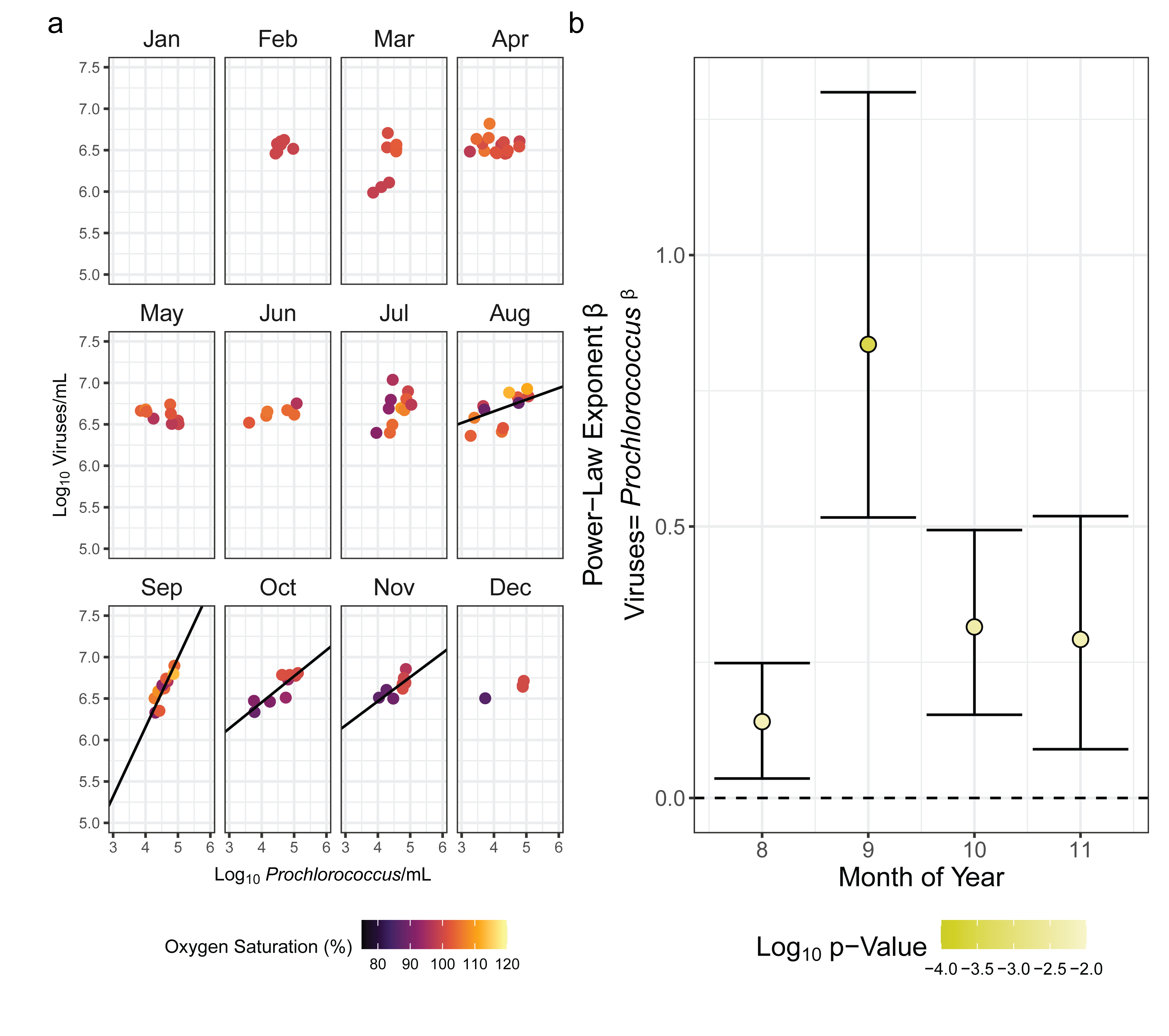

**Supplementary Figure 12.** **Seasonal emergence of power-law relationships between *Prochlorococcus* abundance and bulk virus-like-particle (VLP) abundances across the BATS historical time series.** a) Observations of VLP counts from Parsons *et al*.^3^ with paired *Prochlorococcus* counts from the BATS time series, colored by oxygen saturation calculated from CTD oxygen, temperature, and salinity data. Solid line is major-axis regression best fit of the relationship between log_10_(*Prochlorococcus*) and log_10_(VLP). Major-axis regression lines are only plotted when the slope is significantly different from 0 based on bootstrapped simulations to estimate uncertainty in the slope. b) Power law exponents (linear regression of data plotted in panel a) by month in the form viruses=(intercept)**Prochlorococcus*^slope. Points represent point estimates of the power law exponent (slope of the regression in panel a), and vertical bars represent 95% confidence intervals on the estimate of the slope. The model significance as indicated by fit p-value is the color of the point. Slopes with uncertainties are plotted for models with a significance level of p<0.05.
